# The penetration of sunflower root tissues by the parasitic plant *Orobanche cumana* Wallr. is intracellular

**DOI:** 10.1101/2023.07.24.550254

**Authors:** MC Auriac, C Griffiths, A Robin-Soriano, A Legendre, MC Boniface, S Muños, J Fournier, M Chabaud

## Abstract

Parasitic plants cause yield losses for important crops. Among these, *Orobanche cumana* Wallr, sunflower broomrape, is one of the major pests for sunflower. Previous studies stated that in most cases, the haustorium, a specific parasitic plant organ, penetrates host roots intercellularly. However, host cellular mechanisms involved during the parasitic cells penetration remained poorly described.
We investigated sunflower root cellular behavior during haustorium penetration using various microscopy approaches including live cell imaging of inoculated transgenic fluorescent sunflower roots.
We showed that the haustorium of *O. cumana* penetrated living sunflower root tissues, as a result of the degradation of the host cell wall and the formation of a new host trans-cellular apoplastic compartment for haustorium accommodation. Moreover, broomrape induced cell divisions in outer root tissues at very early stages of the interaction, leading to localized hypertrophy at the site of broomrape attachments.
These findings are a change of paradigm in the research field of parasitic plants. They extend host root intracellular accommodation mechanisms initially shown for symbiotic and pathogenic biotrophic fungi to parasitic plants. It paves the way for future understanding and development of resistance to parasitic plants.

**Key message:** Combination of *in vivo* confocal, large field and transmission electron microscopy approaches revealed how intimate the relationship between the parasitic plant broomrape (*Orobanche cumana* Wallr.) and its sunflower host (*Helianthus annus* L.) is at very early stages of their interaction.

## Introduction

Sunflower broomrape (*Orobanche cumana*) is one of the main pests for sunflower crops. This holo-parasitic plant, is specific to sunflower crops. Broomrape seeds perceive their host thanks to germination stimulants present in sunflower root exudates (Bouwmeester et al., 2021). Once germinated, the broomrape radicle grows toward the host root (Krupp et al. 2021) and develops papillae, which adhere to the host root and secrete mucilaginous compounds (Joel and Losner-Goshen, 1994). Subsequently, epidermal cells at the tip of the haustorium, a specific parasitic organ, differentiate into intrusive cells that penetrate the host root (Masumoto et al., 2021). This penetration combines physical pressure and degradation of sunflower root cell walls thanks to pectolytic activity enzymes released by the parasitic plant (Shomer-Ilan, 1993; Losner-Goshen et al., 1998). Intrusive cells make their way toward the host root vessels, crossing the successive host root tissues. Transcriptomic analyses showed that, in the case of a susceptible interaction, defense genes were activated only transiently and at a low level (Letousey et al., 2007; Dos Santos et al., 2003a, b). In addition, the expression of the putative defence suppressor gene *Par1* of various parasitic plants at the early stages of interaction (Yang et al., 2020; Qiu et al., 2022), suggests manipulation of their host by parasitic plants. Once in contact with the host xylem vessels, intrusive cells differentiate into vessel elements and vascular connections are established (xylem as well as phloem), to insure the nutrient supply of the parasite (Krupp et al., 2019). Although numerous studies have been performed on parasitic seed germination and haustorium development (Yoshida et al. 2016), most of them were focused on the parasitic plants, and the host cellular mechanisms involved during the intrusive cell development were poorly described (Mutuku et al., 2021). How the host cells behave during the massive expansion of the haustorium tissues across the outer root cell layers remained quite unknown. A few studies published in the 70-90s explored the host cellular reorganization during the early stages of the haustorium penetration of various *Orobanchaceae* parasitic plant species. It was shown that haustorium development is accompanied by unusual host cell proliferation (Kuijt 1977; Dörr and Kollmann, 1974). Whether the penetration is intra or intercellular in the root host was rarely stated. Dörr and Kollman (1974) and Kuijt (1977) mentioned intercellular growth only of the haustorial cells, with no observations of plasmodesmata interconnecting host and parasite cells for the interactions *O. crenata/ Vicia faba* and *O. ramosa/ Cannabis sativa*. Intercellular penetration between two cortical cells was shown during the interaction between *Striga gesnerioides* (another *Orobanchaceae* species) and cowpea (*Vigna unguiculata;* Reiss and Bailey, 1998). By contrast, the work by Dörr (1969) on the stem parasitic plant species *Cuscuta* (*Convolvulaceae* family) on the host *Pelargonium zonale* revealed intracellular as well as intercellular penetrations preceding the vascular connection between the host and the parasite (Press et al., 1990). In addition, Musselmann and Dickinson (1975) showed an example of an intrusive cell of the parasitic plant *Agalinis aphylla* (*Orobanchaceae* family) penetrating intracellularly a cortical cell through a small opening in the cell wall. Thus, whether sunflower root penetration by the broomrape haustoria is intra and/or inter cellular remained an open question. This knowledge is required in the perspective of subsequently investigate and understand the sunflower cellular mechanisms associated with resistances to *O. cumana*. In this work, using an efficient selection of the early stages, and combining various microscopy approaches including live-cell imaging of transgenic fluorescent host tissues, we re-investigated the relationships between host and parasitic tissues at the cell level during the early stages of haustorium penetration. The questions we addressed were: *(i)* do intrusive cells penetrate the host root inter or intra-cellularly? *(ii)* do the sunflower root cells in the vicinity of the intrusive cells die or stay alive? *(iii)* are sunflower cell divisions induced at the very early stages of the penetration, and which are the root tissues involved?

### Early stage kinetics of the sunflower/ broomrape interaction in rhizotrons

To answer these questions, a dedicated growth and inoculation device called rhizotron (Le Ru et al., 2021, **Notes S1**) was used in order to select very young attachments, *i*.*e*. haustorium penetration before vessel connections. We used young inoculated wild-type sunflower plantlets (*i*.*e*. non transformed) for longitudinal sections of attachments (large field microscopy and transmission electron microscopy [TEM]). In a second approach, attachments were captured as *in vivo* confocal microscopy images making use of living transgenic composite sunflower plants, expressing the Green Fluorescent Protein (GFP) tagged to the endoplasmic reticulum (ER) (**Fig. S1-S2**; **Table S1**). Observation of attachments was performed from 4 to 8 days after inoculation (dai) (**Table S2**). Broomrape rarely penetrated the host root before 6 dai, while most of the haustoria had reached the inner root tissues (inner cortex to the vessels) at 8 dai. The kinetics were very similar whatever the type of plants and microscopy approach. Interestingly, similarly to our observations, Joel and Losner-Goshen (1994) observed the first stages of attachments at 5-7 dai.

### Broomrape enters into living sunflower root tissues intracellularly

Germinated broomrape seeds developed papillae at the tip of the radicle when contacting the host root (**Fig. S2i**, Joel and Losner-Goshen, 1994). Mechanical pressure of the broomrape in contact with sunflower root epidermal cells led to cell wall deformation (**Fig. 1a-c**). Differentiated intrusive cells at the broomrape radicle tip were strongly stained by toluidine blue O. They displayed a very dense cytoplasm, a reduced vacuole and a large nucleus containing a darkly stained nucleolus, suggesting a high metabolic activity (**Fig.1a, d, g, j**). Imaging early stages of broomrape penetration revealed that intrusive cells penetrated the epidermal layer as well as the successive outer cortical layers intracellularly (**Fig. 1d-i**). In our culture system, sunflower roots had 4 to 5 cortical cell layers between the epidermis and the endodermis (**Fig. S3**). Intracellular penetration of sunflower root cells was observed in all the analyzed penetration sites (21 sites for large field microscopy and 21 sites for confocal microscopy, **Table S2**). The use of the GFP-ER construct provided information about both the cytoplasmic organization and the nucleus position, thanks to the ER outline labelling the nuclear envelope (Genre et al. 2005). In many cases, the nucleus of the penetrated cell was strikingly positioned close to the intrusive cells (**Fig. 1b, c, h, i**), suggesting perception of the intrusive cell, either through the exerted mechanical pressure (Genre et al. 2009) or through unknown chemical signals. However, in contrast to root penetration by symbiotic (Genre et al., 2005; 2008) or pathogenic biotrophic fungi (Koh et al., 2005; Kankanala et al., 2007; Genre et al., 2009), no cytoplasmic aggregation, nor specific ER re-organization were observed ahead of the penetration process. Interestingly, ER was surrounding the broomrape intrusive cells (**Fig. 1e, f, h, i, k, l**), showing active, though not massive, host intracellular re-organization along with the penetration process. These results suggested active membrane synthesis around intrusive cells requiring nucleus and ER activity in the host cell, and showed that the sunflower penetrated cells remained alive. Deeper root tissues (endodermis and pericycle) were also penetrated intracellularly by intrusive cells (**Fig. 1j**). In most cases, haustoria penetrated the host root with minimal host cell damage. However, the live-cell imaging approach revealed a few cases of cell death as shown by the absence of fluorescence (2 sites, **Fig. S4a, b**), or a severe ER disruption (1 site, **Fig. S4c, d**). Similarly, change of the vacuole structure was observed using large field microscopy for a few sites (7 sites among 21 penetration sites): appearing as a blue smear (**Fig. 1d; Fig. S4e**) or light blue material filling the cell (**Fig. S4e**). One or a few penetrated cells only were affected, adjacent to the intrusive cells in outer root tissues. These results suggested that in some cases, penetration of the intrusive cells got out of control and synchronization of the penetration process and the sunflower cellular reorganization failed, leading to sunflower cell death. This phenomenon remained cell-autonomous, without other defense reactions in the surrounding or the deepest root tissues. Furthermore, penetration could result in the separation of the host nucleus from the distal part of the penetrated cell, probably leading to cell death as well. Strikingly, broomrape intrusion was thicker in outer root tissues (**Fig. 1)** than in inner root tissues, in which only single elongated and separated intrusive cells were detected (**Fig. 1e, f, h, i, k, l**). Similarly, Dörr et al. (1969) reported intracellular “searching hyphae” for the *Cuscuta* stem parasite. Sunflower root cell divisions were observed as early as 6 dai, close to attachments (respectively 9 and 7 sites for sections and live-cell imaging). Divisions were mostly anticlinal in the cortex, and periclinal in the pericycle (**Fig. S5; Fig. 1d, e, h, i**). The number of dividing root cell layers and the length of the dividing zone were highly variable (for example from 1 cortical cell to more than 30 cells in a row). These divisions may account for root hypertrophy that was previously observed at the site of 14-dai attachments in rhizotrons (Chabaud et al., 2022). Sunflower roots were known to swell locally at the site of broomrape attachment by means of cell division (Kuijt, 1977) and as early as 7 dai (Dörr and Kollamnn, 1994). These divisions could be induced indirectly (host hormonal regulation) or directly by the parasitic plant (hormonal release: such as auxin [Ishida et al., 2016] or cytokinin [Spallek et al., 2017]). Whether germinated broomrape seed exudates would be sufficient for the induction of host cell divisions remains an open question.

**Fig. 1.**
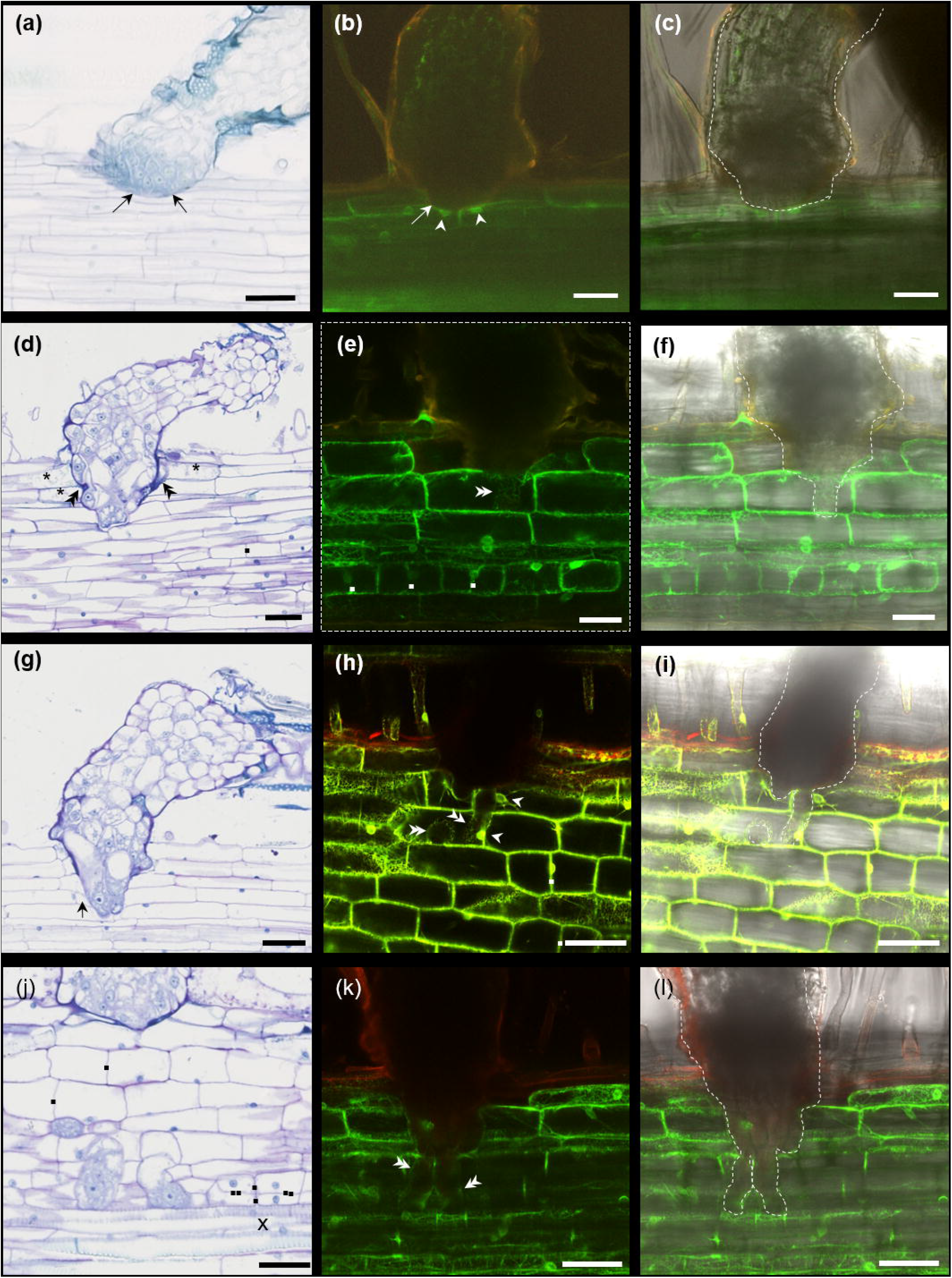
Broomrape haustorium penetrates the sunflower root tissues intracellularly. Broomrape attachments (6-7 dai) were observed at the cellular level, using 2 approaches: *(i)* large field microscopy of thin sections of resin-embedded and toluidine blue O-stained attachments (right column, **a, d, g, j)**; *(ii)* confocal microscopy of broomrape-inoculated transgenic fluorescent sunflower roots expressing the GFP targeted to the endoplasmic reticulum (ER) (middle and left columns, **b, c, e, f, h, i, k, l**). Confocal microscopy images show fluorescence alone (combined GFP fluorescence and auto-fluorescence channels, middle column, **b-e-h-k**) or fluorescence with the corresponding bright field image (left column, **c, f, i, l**). **a-c**. Deformation of the epidermal cell wall (arrow) due to the mechanical pressure exerted by the haustorium. Re-positioning of the nuclei of the deformed epidermal cells in the vicinity of the haustorium tip (**b**. white arrowheads). **d–i** and **k-l**. Intrusive cells penetrated the root epidermal and cortical layers intracellularly. The penetrated cortical cells had wavy (**d**. black double arrowheads) or deformed (**g**. black arrow) cell wall in contact with the haustorium. ER surrounded the intrusive haustorium tip (**h, k**. white double arrowheads). Nuclei were positioned close to the intrusive cells (**h**. white arrowheads). **j**. Intrusive cells reached the host xylem vessels (xylem: black cross), and crossed intracellularly the deepest cortical cell layer as well as the endodermis and the pericycle. Divisions in the cortex (mostly anticlinal, squares) and the pericycle (mostly periclinal, double squares) in the vicinity of the haustorium (**d, e, j). c, f, i, l**. White dots outline the broomape haustorium visible on the bright field image. **b-c, e-f**, **k-l** are z axis projections of serial optical sections. Scale bar = 50 µm (**a, b, c, d, e, f, g, j**); 20 µm (**h, i, k, l**).

### Intrusive cells penetrate a new apoplastic compartment

The interface between intrusive cells and the sunflower penetrated root cells at early stages of the interaction was further characterized by TEM (**Fig. 2)**. A 7 dai attachment with the haustorium reaching the 3^rd^ cortical cell layer is illustrated **Fig. 2a, b**. Starch grains, a sign of the transition from the autonomous (germination stage) to the parasitical stage (Joel and Losner-Goshen, 1993), were present in the central part of the attachment (**Fig. 2a, c**). In the outer root cell layers, the interface appears as a thick layer surrounding broomrape (**Fig. 2b**), as already described for *Striga* (Reiss and Bailey, 1998; Neuman et al., 1999). The intrusive cells were easily distinguished from sunflower root cortical cells thanks to their dense cytoplasm, containing Golgi stacks, large mitochondria as well as a reduced vacuole (**Fig. 2b, e, f**) as already reported by Kuijt and Toth (1976) and Kuijt (1977). By contrast, the host cortical cells whether penetrated or not, contained a large vacuole with a thin layer of surrounding cytoplasm (**Fig. 2b, d**). Mitochondria in the penetrated host cells were present all along the host plasmalemma, suggesting intense activity at the periphery of the host cell such as membrane biosynthesis (**Fig. 2d)**. This dense cytoplasm confirmed that penetrated host cells were alive at this stage. The parasitic cell wall was present all around the intrusive cells. By contrast the host cell wall on the anticlinal interface seemed to be missing (**Fig. 2e, f**) or disorganized **(Fig. 2d, g, h**). At the frontline of the haustorium penetration the disorganization of the host cell wall was less pronounced (**Fig. 2i**), suggesting progressive local enzymatic degradation of the host cell wall. This apparent dissolution of the nearby host cell walls (Kuijt, 1977) or, in the case of the interaction *Ptheirospermum japonicum*/ *Arabidopsis thaliana*, the cell wall partial digestion at the interface (Kurotani et al., 2020, for the interaction) had been reported previously. Neverthless, the host cell plasmalemma seemed to remain undisturbed and continuous (**Fig. 2g; Fig. S6b**). Both host and parasitic plasma membranes were highly convoluted at the front line of the haustorium (**Fig. S6b)**, suggesting membrane synthesis for the haustorium accommodation (host) and haustorium expansion (parasite). No plasmodesmata were observed on the interface at these early stages, indicating that molecular exchanges between the parasite and the host happened at later stages, or through vessel connections, as interspecific plasmodesmata have been shown in the phloem (Krupp et al., 2019). In some cases, the penetration led to disaggregation of the vacuole of the host cell (**Fig. S6c-d**), with disruption of the host cell plasmalemma, leading to cell necrosis, as for *Striga* (Neuman et al., 1999). However as mentioned above this was not very common and remained cell autonomous. In addition, no evidence of cell death was observed at the later stages (Chabaud et al., 2022). Altogether, these results showed that the parasitic intrusive cells penetrate the host root cells intracellularly, as a result of degradation of the host cell wall and formation of a new host trans-cellular apoplastic compartment for haustorium accommodation.

**Fig. 2.**
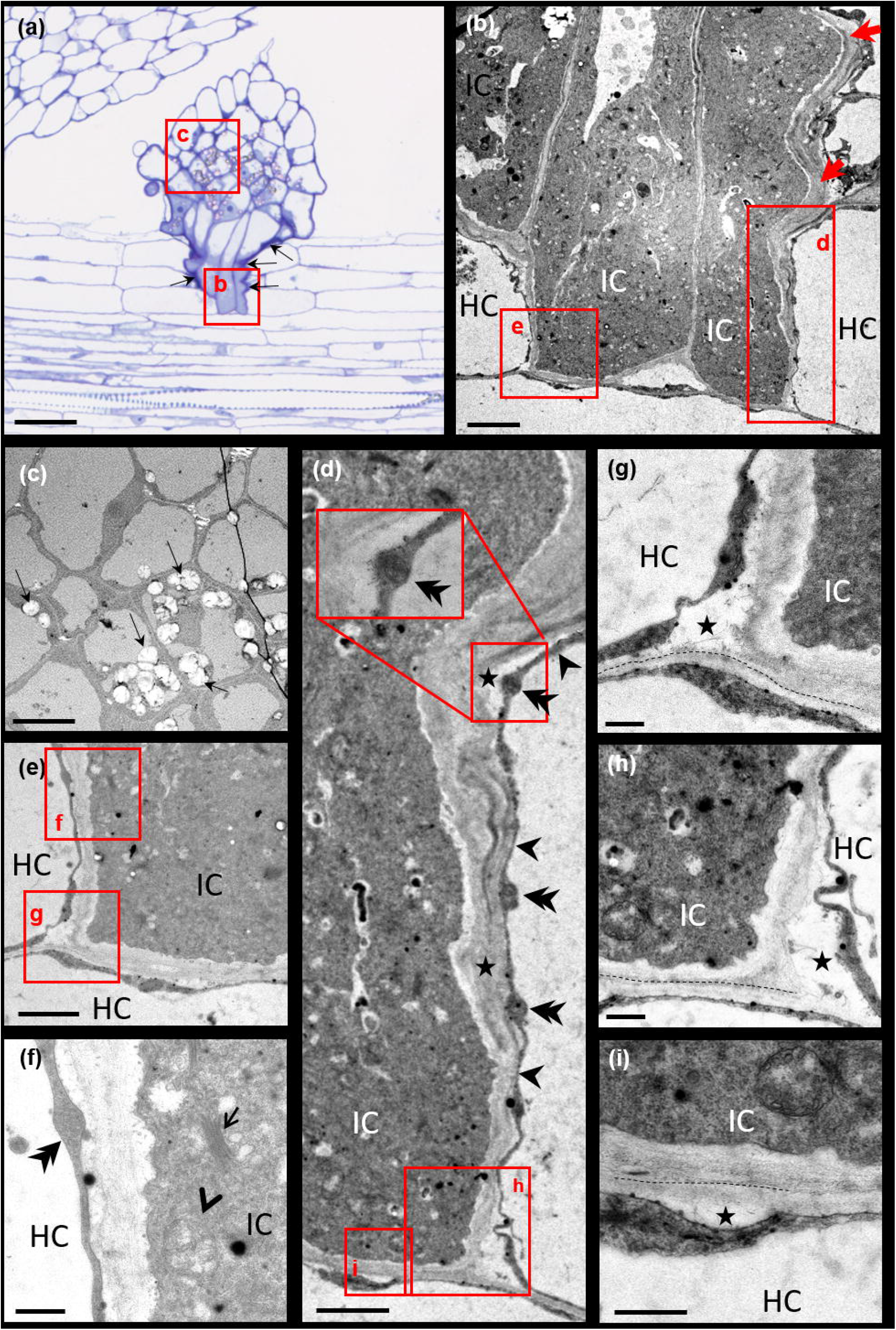
Creation of a new apoplastic compartment for haustorium accommodation. A 7 dai attachment, with intrusive cells reaching the 3^rd^ cortical cell layer was imaged. **a**. General view of the attachment using large field microscopy. **b-i**. Transmission electron microscopy. IC intrusive cell, HC host cell. **a, b**. Intrusive cells intracellular penetration in the host outer cortex cell. A thick stained layer surrounded the penetrating intrusive cells in outer host tissues (epidermis and outer cortex; black arrows in **a**, red arrows in **b**). **c**. Magnification of the central part of the broomrape, with starch grain-containing cells. **d**. Disorganization of the host cell wall (star) on the anticlinal interface between the intrusive cell and the penetrated host outer cortical cell. Continuous thin layer of cytoplasm in the host cortical cell (black arrowheads), containing numerous mitochondria (double arrowheads). **e, f**. Presence of the parasitic cell wall but not the host cell wall on the anticlinal interface. **e**. Magnification of **b. f**. Magnification of **e. f**. The intrusive cell cytoplasm contains Golgi (thin arrow) and mitochondria (arrowhead). Mitochondria are also present in the thin host cytoplasm layer (double arrowhead). **g-h**. At the tip of the haustorium, complete disorganization of the host cell wall (star) with presence of fibrillae fragments. Convolution of the plasmalemma of the host cortical cells adjacent to the intrusive cells. **i**. At the front line of the intrusive cell tips (periclinal interface), local disorganization of the host cell wall (star), the plasmalemma of the host cell remaining intact and continuous. The dotted line shows the separation between the parasitic and host cell walls (**g-i**). Scale bar = 50 µm (**a**); 5 µm (**b**); 10 µm (**c**); 2 µm (**d-e**); 0.5 µm (**f, g, h, i**).

### Concluding remarks

Most striking among our findings has been the observation of intracellular haustorium penetration of host root tissues, in contrast to most studies on *Orobanchaceae*. These studies relied mainly on the observation of transverse sections, by contrast to the longitudinal sections used in this work, which made it easier to distinguish between intra and intercellular processes. Our work showed the intimate broomrape penetration into its host, through the formation of a new apoplastic compartment. It suggested that although host cell wall integrity has been damaged by parasitic cell wall degrading enzymes, only minor defense reactions were induced as previously reported for biotrophic pathogenic fungi (Mendgen and Hahn, 2002; Bellincampi et al., 2014). In that respect, genes encoding cell wall degrading enzyme inhibitors could be good candidates for resistance to broomrape. In addition, as *HaOr7* (Duriez et al. 2019) and *HaOr*_*Deb2*_ (Fernandez-Aparicio et al., 2022) encode Leucine-Rich-Repeat Receptors Like Proteins, providing resistance to various *O. cumana* races, it would be of outstanding interest to characterize the cellular processes involved in these incompatible interactions. Comparing the cellular processes for various *O. cumana* races could highlight common or different mechanisms. Finally, using these approaches on other major parasitic plant species such as *Striga* will be of great interest for future resistance development in a larger host range.

## Acknowledgments

We thank P. Gresshoff (University of Queensland, Australia) for the *A. rhizogenes* strain K599. This study was supported by the “Laboratoires d’Excellences (LABEX)” “Towards a Unified theory of biotic interactions: roLe of environmental Pertubations” (TULIP; ANR-10-LABX-41)” and/or by the “École Universitaire de Recherche (EUR)” TULIP-GS (ANR-18-EURE-0019). Slide scanning was performed using the Nanozoomer from the Imagery Platform of the Federated Research Institute AgroBiosciences-Interactions-Biodiversité (FRAIB; Castanet-Tolosan, France). This study was performed in the frame of a 2-year project (SunOrCell), funded by “Promosol/ SeleoPro” (the association of French Sunflower and Rapeseed Breeders for promoting these crops). The International Consortium of Sunflower Genomics (ICSG) supported the grants for the training students CG and ARS.

## Authors and contribution

MCA performed cytological experiments (large field microscopy and TEM). CG, ARS and AL established sunflower transformation experiments. MCB produced sunflower and broomrape resources. SM: coordinated the ICSG, assisted with the construction of the project and the writing of the manuscript. JF gave technical and scientific advice, assisted with the construction and the writing of the manuscript. MC designed the experiments, carried out confocal microscopy and cytology experiments, wrote the manuscript.

## Data availability statement

The original contributions presented in the study are included in the article/Supplementary Material. Further inquiries can be directed to the corresponding author.

## References

Bellincampi D, Cervone F, Lionetti V. 2014. Plant cell wall dynamics and wall-related susceptibility in plantpathogen interactions. Frontiers in Plant Science 5: 228.

BouwmeesterH, Li C, Thiombiano B, Rahmini M, Dong L. 2021. Adaptation of the parasitic plant lifecycle: germination is controlled by essential host signaling molecules. Plant Physiology 185: 1292–1308.

Chabaud M, Auriac M-C, Boniface M-C, Delgrange S, Folletti T, Jardinaud M-F, Legendre A, Perez-Vich B, Pouvreau J-B, Velasco L, Delavault P, Muños S. 2022. Wild Helianthus species: A reservoir of resistance genes for sustainable pyramidal resistance to broomrape in sunflower. Frontiers in Plant Science 13.

Dörr I. 1969. Feinstruktur intrazellular wachsender Cuscuta-Hyphen? Protoplasma 67: 123–137.

Dörr I, Kollmann R. 1974. Strukturelle Grundlage des Parasitismus bei Orobanche I. Wachstum der Haustorialzellen im Wirtsgewebe. Protoplasma 80: 245–279.

Dörr I. 1997. How Striga parasitizes its host: a TEM and SEM study. Ann. Bot. 79:463–72.

Dos Santos CV, Delavault P, Letousey P, Thalouarn P. 2003a. Identification by suppression subtractive hybridization and expression analysis of Arabidopsis thaliana putative defence genes during Orobanche ramosa infection Physiology Molecular Plant Pathoogyl 62: 297–303.

Dos Santos CV, Letousey P, Delavault P, Thalouarn P. 2003b. Defense Gene Expression Analysis of Arabidopsis thaliana Parasitized by Orobanche ramosa. Phytopathology 93: 451–457.

Duriez P, Vautrin S, Auriac MC, Bazerque J, Boniface MC, Callot C, Cauet S, Chabaud M, Gentoux F, Lopez-Sendon M et al. 2019. A receptor-like kinase enhances sunflower resistance to Orobanche cumana. Nature Plants 5: 1211–1215.

Fernandez-Aparicio M, del Moral L, Muños S, Velasco L, Pérez-Vich B. 2022. Genetic and physiological characterization of sunfower resistance provided by the wild-derived Or_Deb2_gene against highly virulent races of Orobanche cumana Wallr. Theoritical Applied Genetics 135 : 501–525

Genre A, Chabaud M, Timmers ACJ, Bonfante P, Barker DG. 2005. Arbuscular mycorrhizal fungi elicit a novel intracellular apparatus in Medicago truncatula root epidermal cells before infection. Plant Cell 17: 3489–3499.

Genre A, Chabaud M, Faccio A, Barker DG, Bonfante P. 2008. Prepenetration apparatus assembly precedes and predicts the colonization patterns of arbuscular mycorrhizal fungi within the root cortex of both Medicago truncatula and Daucus carota. Plant Cell 20: 1407–1420.

Genre A, Ortu G, Bertoldo C, Martino E, Bonfante P. 2009. Biotic and abiotic stimulation of root epidermal cells reveals common and specific responses to arbuscular mycorrhizal fungi. Plant Physiology 149: 1424–1434.

Ishida JK, Wakatake T, Yoshida S, Takebayashi Y, Kashara H, Wafula E, dePamphilis CW, Namba S, Shirasu K. 2016. Local auxin biosynthesis mediated by a YUCCA flavin monooxygenase regulates haustorium development in the parasitic plant Phtheirospermum japonicum. Plant Cell 28: 1795–1814.

Joel DM, Losner-Goshen D. 1994. The attachment organ of the parasitic angiosperms Orobanche cumana and O. aegyptiaca and its development. Canadian Journal Botany 72: 564–574.

Kankanala P, Czymmek K, Valent B. 2007. Roles for rice membrane dynamics and plasmodesmata during biotrophic invasion by the blast fungus. Plant Cell. 19: 706–724.

Koh S, Andrea A, Edwards H, Ehrhardt D, Somerville S. 2005. Arabidopsis thalinan subcellular responses to compatible Erysiphe cichoracearum infections. Plant Journal 44:516–529.

Krupp A, Heller A, Spring O. 2019. Development of phloem connexion between the parasitic plant Orobanche cumana and its host sunflower. Protoplasma 256: 1385–1397.

Krupp A, Bertsch B, Spring O. 2021. Costunolide Influences Germ Tube Orientation in Sunflower Broomrape A First Step Toward Understanding Chemotropism. Frontiers in Plant Science 12.

Kuijt J. 1977. Haustoria of phanerogamic parasites. Annual Review Phytopathology 17: 91–118.

Kuijt J, Toth R. 1976. Ultrastructure of angiosperm haustoria-A review. Annals Botany 40: 1121–30.

Kurotani KI, Wakatake T, Ichihashi Y, Okayasu K, Sawai Y, ogawa S, Cui S, Suzuki T, Shirasu K, notaguchi M. 2020. Host-parasite tissue adhesion by a secreted type of beta-1,4-glucanase in the parasitic plant Phtheirospermum japonicum. Communications Biology 3.

Le Ru A, Ibarcq G, Boniface MC, Baussart A, Munos S, Chabaud M. 2021. Image analysis for the automatic phenotyping of Orobanche cumana tubercles on sunflower roots. Plant Methods 17.

Letousey P, de Zelicourt A, Dos Santos CV, Thoiron S, Monteau F, Simier P, Thalouarn P, Delavault P. 2007. Molecular analysis of resistance mechanisms to Orobanche cumana in sunflower. Plant Pathology 56: 536–546.

Losner-Goshen D, Protnoy VH, Mayer AM, Joel DM. 1998. Pectolytic Activity by the Haustorium of the Parasitic Plant Orobanche L.(Orobanchaceae) in Host Roots. Annals Botany 81: 319–326.

Masumoto N, Suzuki Y, Cui S, Wakazaki M, Sato M, Kumaishi K, Shibata A, Furuta K, Ichihashi Y, Shirasu K et al. 2021. Three-dimensional reconstructions of haustoria in two parasitic plant species in the Orobanchaceae. Plant Physiology 185: 1429–1442.

Mendgen K, Hahn M. 2002. Plant infection and the establishment of fungal biotrophy. Trends in Plant Science 7: 352–356.

Mutuku JM, Cui S, Yoshida S, Shirasu K. 2021. Orobanchaceae parasite–host interactions. New Phytologist 230: 46–59.

Musselman LJ, Dickison WC. 1975. The structure and development of the haustorium in parasitic Scrophulariaceae. Botanical Journal Linn. Society 70: 183–212.

Neumann U, vian B, weber HC, Sallé G. 1999. Interface between haustoria of parasitic members of the Scrophulariaceae and their hosts: a histochemical and immunocytochemical approach. Protoplasma 207: 84–97.

Press MC, Gravest JD, Stewart GR. 1990. Physiology of the interaction of angiosperm parasites and their higher plant hosts. Plant Cell and Environment 13: 91–104.

Qiu S,Bradley JM, Zhang P, Chaudhuri R, Blaxter M, Butlin RK, Scholes JD. 2022. Genome-enabled discovery of candidate virulence loci in Striga hermonthica, a devastating parasite of African cereal crops. New Phytologist 236: 622–638.

Reiss GC, Bailey JA. 1998. Striga genesrioides parasiting cowpea: development of infection structures and mechanisms of penetration. Annals Botant 81: 431–440.

Shomer-Ilan A. 1993. Germinating seeds of the root parasite Orobanche aegyptiaca Pers. excrete enzymes with carbohydrase activity. Symbiosis 15: 61–70.

Spallek T, Melnyk CW, Wakatake T, Zhang J, Sakamoto Y, Kiba T, Yoshida S, Matsunaga S, Sakakibara H, Shirasu K. 2017. Interspecies hormonal control of host root morphology by parasitic plants. Proceedings National Academy Science USA 114: 5283–5288.

Yang C, Fu F, Zhang N, Wang J, Hu L, Islam F, Bai Q, Yun X, Zhou W. 2020. Transcriptional profiling of underground interaction of two contrasting sunflower cultivars with the root parasitic weed Orobanche cumana. Plant Soil 450: 303–321.

Yoshida S, Cui S, Ichihashi Y, Shirasu K. 2016. The Haustorium, a Specialized Invasive Organ in Parasitic Plants. Annual Review Plant Biology 67:643–67.

