## Supplementary material for "The penetration of sunflower root tissues by the parasitic plant *Orobanche cumana* Wallr. is intracellular": Notes S1; Fig. S1; Fig. S2; Fig. S3; Fig S4; Fig. S5; Fig. S6; Table S1; Table S2

**Notes S1. Details of the materials and methods.**

***Plant genotypes, bacterial strains, and constructs***

We used the susceptible cultivated *H. annuus* L., XRQ (Chabaud et al., 2022; Badouin et al., 2017) as the sunflower host for experiments in rhizotrons and for root transformation experiments. The *O. cumana* Wallr. populations used in this study was the French race E-BOU of *O. cumana,* with a virulence level classified between E and F, and harvested in 2017 in Bourret (Tarn et Garonne, France; reference: LIPM-20734).

The *Agrobacterium rhizogenes* strain K599 (Savka et al. 1990) was used for sunflower root transformation. The binary vector used was derived from pBIN19 (Bevan 1984) and carried the construct p35S-GFP-ER coding for the Cauliflower Mosaic Virus 35S promoter (p35S) driving the constitutive expression of the Green Fluorescent Protein tagged to the endoplasmic reticulum due to the presence of ER-targeting sequences (N-terminal signal peptide) and ER-retention (C-terminal tetra-peptide HDEL; Haseloff et al. 1997). Bacteria were grown on LB medium supplemented with Streptomycin 100 mg/l (bacterial selection) and 50 mg/l (plasmid selection)

***Experiment in rhizotrons for studying the early stages of the interaction between non-transformed sunflower plants and broomrape***

Sunflower seeds of wild-type (*i.e.* non-transformed) plants were surface sterilized for 10 min in a 4.8 % sodium hypochlorite solution, rinsed 3 times and sown in a 1/1 v/v mixture of sand/ vermiculite in alveoli. Seedlings were grown at 22 °C, 60% humidity, 16 h light 118 µE/m²/s for 7 days. Broomrape seeds were surface sterilized for 5 min in a 3.2 % sodium hypochlorite solution, rinsed 3 times using a 40 µm cell strainer, and water-conditioned for 7 days in a 50 ml sterile tube at 23 °C in the dark, at a final concentration of 10 mg/ 3 ml of water. Rhizotrons, 12x12 cm home-made plexiglass boxes were assembled as described in Le Ru et al. (2021), with water-soaked sterile rock-wool, a sterile glass fiber paper and a 7 day-old sunflower plantlet. Each rhizotron was inoculated with 10 mg (3 ml) of conditioned broomrape seeds.

***Large field microscopy observations***

This approach was inspired from the work of Xiao et al. (2014) on young symbiotic nodule development. Four to 8 dai, root samples were prepared for cytological studies as described in Chabaud et al. (2022). Fixed samples were embedded in Technovit 7100 resin (Heraeus Kulzer,Wehrheim, Germany), according to the manufacturer’s recommendations. Thin (4-5 µm) sections, longitudinal to both haustorium and host root, were made using a microtome (2040 Reichert Jung), stained in 0.2% toluidine blue for 3 min, mounted in DePeX mounting medium (BDH Laboratories, Poole, England) and scanned using a Nanozoomer (NDP, Hamamatsu).

***Transmission electron microscopy (TEM) observations***

Samples were prepared as in Cerutti et al. (2017) with some modifications. Five to 7 dai broomrape attachments on sunflower root fragments were fixed under vacuum for 30 min with 2.5 % glutaraldehyde in 0.1 M sodium cacodylate buffer (pH = 7.2) containing 0.1 % Triton X100, and then at the atmospheric pressure in the same solution without Triton X100, washed in the same cacodylate buffer and post-fixed for one hour at room temperature with 1 % osmium tetroxide ($OsO_4$) in the same buffer. They were then dehydrated in ethanol series, and embedded in Epon. Thin (1 µm) or ultra-thin (80-90 nm) sections were prepared on UltraCut E ultramicrotome (Reichert-Jung) equipped with a diamond knife (Reichert-Leica, Germany). The histological organization of tissues was observed on thin sections stained in a 1 % borax solution containing 0.2 % methylene blue and 0.1 % toluidine blue, rinsed in water and then in an aqueous solution of 0.07 % basic fuchsin. Ultrastructural observation was done on ultrathin sections stained with uranyless (Delta microscopy, Mauressac France), and lead citrate (delta microscopies, Mauressac France) using the electron microscope Hitachi-7700 (Japan) operating at 80 kV.

***Sunflower root transformation by* A. rhizogenes**

To transform sunflower roots and obtain composite plants, we used a modified version of the protocol from Parks and Yordanov (2020, **Fig. S1-S2**). Sunflower seeds were decontamined with 4.8% sodium hypochlorite for 10 min, rinsed three times with sterile water, let for 1 hour in water before sowing in seedling trays previously filled with a watered mixture of 1/1 (v/v) vermiculite and sand. Seedlings were grown at 22 °C, 60% humidity, 16 h light 118 µE/m²/s for 10 days. Plantlets were watered with a nutrient solution ½ Long Ashton supplemented with 370 µM of phosphate. Transformation of plantlets was performed according to Parks and Yordanov in soaked pre-cut rock-wool cubes (reference ALR02G from GRODAN) with bacterial solution [final OD= 0.25 in ¼ (MS + Gamborg vitamins B5)], in Magenta boxes. After 3 days, Magenta boxes were slightly opened. Six days after transformation, plantlets were transferred to hydroponics, using sterile 1000µl cone boxes, as described in Morel et al. (2018) for another 6 days of culture in ¼ MS liquid medium (without vitamins; Sigma reference MS

5524). Culture in hydroponics was very beneficial to newly developed root growth. Finally, 14 to 18 days following transformation (6-7 days in rock-wool cubes and 7-12 days in hydroponics; **Fig. S1**), transformation efficiency was recorded by measuring the percentage of plants with fluorescent roots and the number of fluorescent roots/ transformed plant (**Table S1**). Fluorescence expression was observed by epifluorescence microscopy using a stereomicroscope (Axiozoom V16; Zeiss), equipped with a GFP Long Pass filter (excitation 485/12 nm and emission from 515 LP). Composite plants were transferred to rhizotrons (see below).

Four experiments were performed. In the first experiment, 17-day-old plantlets were used for transformation, and the transfer to rhizotrons was done 18 days later, resulting in large 35-day-old plants that were difficult to handle under the confocal microscope. Hence the length of culture was progressively reduced in the following experiments to 10-day-old plantlets for transformation, culminating with transfer to rhizotrons as soon as 14 days later in experiment No. 4 (**Fig. S1**). The reduction of the age of the plants used for transformation facilitated the manipulation of the composite plants when they were removed from the rock-wool cubes and limited wounding of the transgenic roots. It also facilitated their transfer into rhizotrons and use in confocal microscopy.

***Use of transformed composite sunflower plants in rhizotrons for the observation of the early stages of the interaction between sunflower and broomrape by confocal microscopy***

This approach was inspired from the cytology work of Genre et al. (2005) for arbuscular mycorrhizal symbiosis studies on *Medicago* and carrot. In the case of sunflower composite plants, experiments in rhizotrons were conducted as described above. Instead of 7-day-old plantlets, sunflower composite plants (14 days after transformation) were transferred to rhizotrons. One to 7 days after transfer, sunflower composite plants were inoculated with conditioned and sterilized broomrape seeds (10 mg/ rhizotron). Six days after inoculation (dai), inoculated plants were transferred to a 12x12 cm square Petri dish containing 80 ml of solid medium ½ Long Ashton 370 µM phosphate, 3 g/l Phytagel, with the aerial part outside of the dish (**Fig. S2g**). The root system was covered with a gas-permeable plastic film (Lumox Film, Starsted) as described in Fournier et al. (2015). Attachments were imaged with a Leica TCS SP8 AOBS confocal laser scanning microscope equipped with a long-distance 25X HC FLUOTAR (numerical aperture, 0.95) water immersion objective. The 488 nm argon laser line was used to excite GFP and auto-fluorescence. Specific emission windows used for GFP and auto-fluorescence signals were 500 to 550 nm, and 580 to 650 nm, respectively, and emitted fluorescence was false-coloured in green (GFP), and red (auto-fluorescence). The images shown are single confocal sections or maximal projections of selected planes of a z-stack. Images were acquired and projected using Leica confocal software and processed using Leica confocal software.

### References


**Badouin H, Gouzy J, Grassa, CJ, Murat F, Staton SE, Cottret L, Lelandais-Briere C, Owens GL, Carrere S, Mayjonade B et al. 2017.** The sunflower genome provides insights into oil metabolism, flowering and Asterid evolution. *Nature* **546**: 148

**Bevan M. 1984.** Binary Agrobacterium vectors for plant transformation. *Nuc. Ac. Res* 12:8711–8721

**Cerutti A, Jauneau A, Auriac MC, Lauber E, Martinez Y, Chiarenza S, Leonhardt N, Berthomé R, Noël LD. 2017.** Immunity at cauliflower hydathodes controls systemic infection by *Xanthomonas campestris pv campestris*. *Plant Physiology* **174**: 700–716.

**Chabaud M, Auriac MC , Boniface Mc , Delgrange S , Folletti T , Jardinaud MF , Legendre A, Perez-Vich B, Pouvreau JB , Velasco L , Delavault P, Muños S. 2022.** Wild *Helianthus* species: A reservoir of resistance genes for sustainable pyramidal resistance to broomrape in sunflower. *Frontiers in Plant Science* **13**.

**Fournier J, Teillet A, Chabaud M, Ivanov S, Genre A, Limpens E, de Carvalho-Niebel F, Barker B. 2015**. Remodeling of the Infection Chamber before Infection Thread Formation Reveals a Two-Step Mechanism for Rhizobial Entry into the Host Legume Root Hair. *Plant Physiology* **167** : 1233–1242.

**Genre A, Chabaud M, Timmers ACJ, Bonfante P, Barker DG. 2005**. Arbuscular mycorrhizal fungi elicit a novel intracellular apparatus in *Medicago truncatula* root epidermal cells before infection. *Plant Cell* **17**: 3489-3499.

**Haseloff J, Siemering KR, Prasher DC, Hodge S. 1997**. Removal of a cryptic intron and subcellular localization of green fluorescent protein are required to mark transgenic Arabidopsis plants brightly. *Proceedings National Academy Science USA* **94**:2122–2127.

**Le Ru A, Ibarcq G, Boniface MC, Baussart A, Munos S, Chabaud M. 2021.** Image analysis for the automatic phenotyping of Orobanche cumana tubercles on sunflower roots. *Plant Methods* **17**.

**Morel A, Peeters N, Vailleau, F, Barberis P, Jiang G, Berthomé R, Guidot A . 2018**. Plant pathogenicity phenotyping of *Ralstonia solanacearum* strains. In: Medina C Lopez-Baena FJ, eds. *Host-Pathogen interactions: Methods and Protocols*. **1734**: 223-239.

**Parks T, Yordanov YS. 2020**. Composite plants for a composite plant: An efficient protocol for root studies in the sunflower using composite plants approach. *Plant Cell Tissue and Organ Culture* **140**: 647-659.

**Savka M., Ravillion B., Noe, G., Farrand S.1990**. Induction of hairy roots on cultivated soybean genotypes and their use to prop-agate the soybean cyst nematode. *Phytopathology*.**80**: 503–508.

**Xiao TT, Schilderink S, Moling S, Deinum EE, Kondorosi E, Franssen H, Kulikova O, Niebel A, Bisseling T. 2014.** Fate map of *Medicago truncatula* root nodules. *Development* **141**: 3517-3528.

**Fig. S1. Sunflower transformation experiments and broomrape inoculation of composite plants for confocal microscopy observations.**

The initial experiment (No.1) led to broomrape inoculation of 36-day-old sunflower plants, which were very developed and not easy to handle under confocal microscopy. Hence, the length of culture was progressively reduced in the following experiments to finally reach broomrape inoculation of smaller, 26-day-old plantlets in experiment No.4. This also allowed to generate two series of inoculated plants according to their development in experiments No. 2 and 4, using the most developed composite plants for the 1st inoculation and the less developed composite plants for a 2nd, delayed inoculation 6 days later. Confocal microscopy was performed 5 to 8 dai.

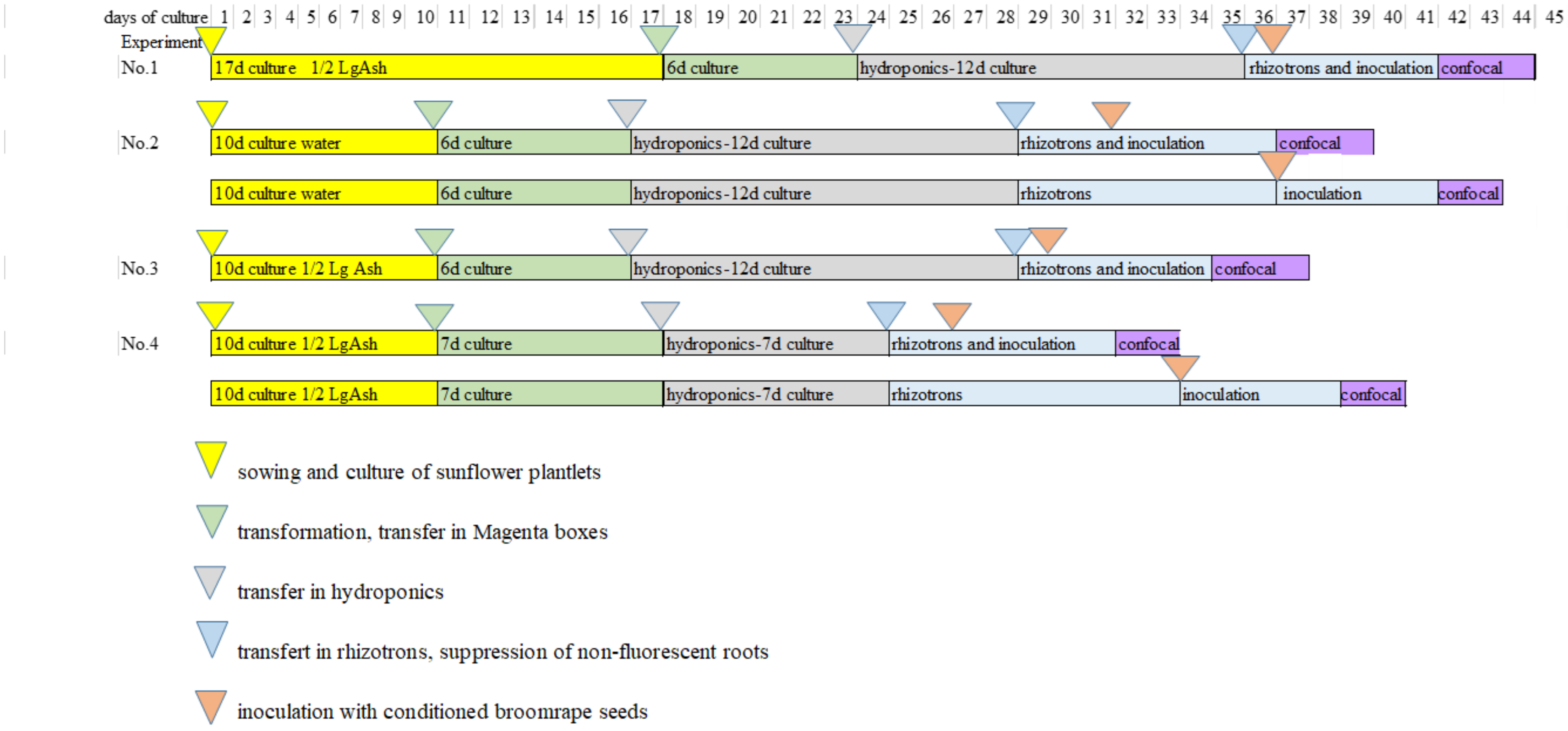

**Fig. S2. Transformation of sunflower plants *via A. rhizogenes* and transfer of composite plants in rhizotrons for broomrape inoculation and observation using confocal microscopy.**

Cuttings without cotyledons were excised from 10-day-old plantlets (**a**), and transformed in pre-soaked rock-wool cubes with *A. rhizogenes* solution (**b**). Six days later, plantlets were transferred in hydroponics (**c**) for another 8 days of culture. Fourteen days after transformation, green fluorescent transgenic roots were counted under a binocular microscope (**d, Table S1**). Non-fluorescent roots were removed and composite plants were transferred to rhizotrons (**e**) and inoculated with pre-conditioned broomrape seeds 1 to 2 days later (**f**). Five to 8 days after inoculation, inoculated composite plants were transferred to a Petri dish (**g-** 5 dai) and attachments were selected and imaged by confocal microscopy (**h-i**). **i.** Papillae development (arrows). Scale bar = 4 cm (**a**); 2 cm (**b**); 5 cm (**c**); 0.5 cm (**d**); 3 cm (**e-g**); 100 µm (**h**); 10 µm (**i**).

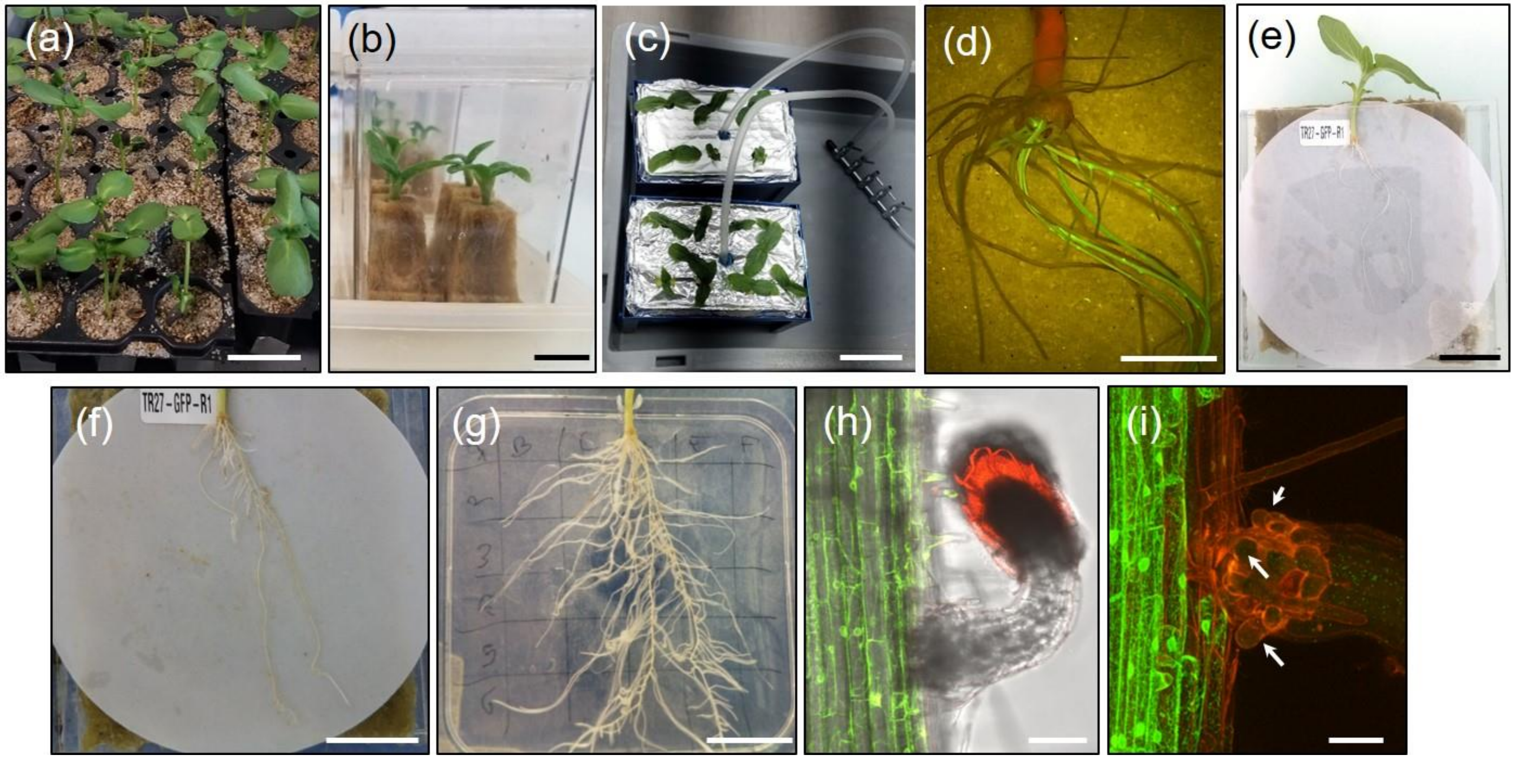

**Fig. S3. Longitudinal section of a sunflower root.**

A thin section of a sunflower root fragment, without broomrape attachment, resin-embedded and stained with toluidine blue O, was observed using large field microscopy. In our growth conditions, roots displayed 4 to 5 cortical layers.

Scale bar = 100 µm.

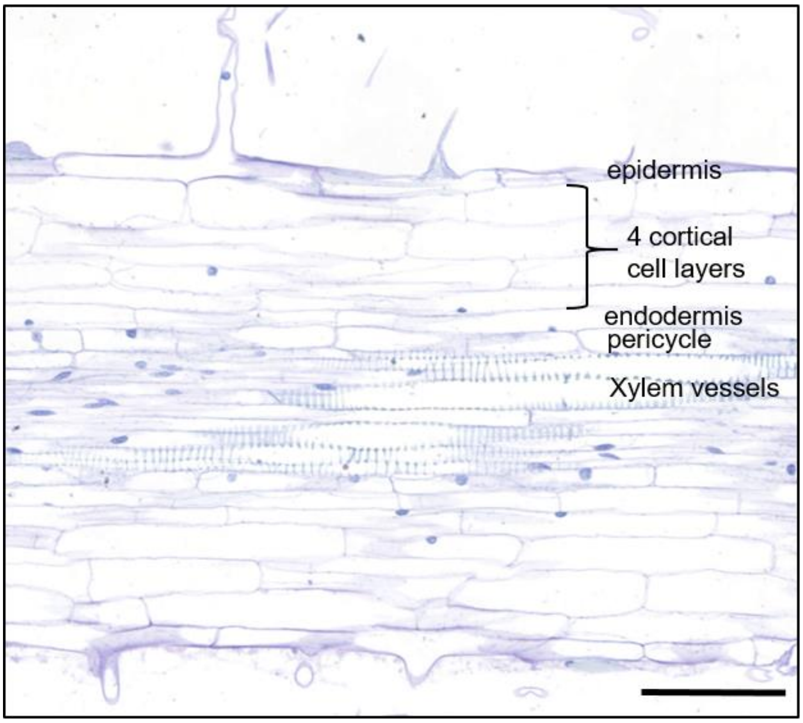

**Fig. S4. Cell death or ER de-structuring associated with haustorium penetration.**

Confocal microscopy imaging revealed that occasionally, sunflower cells in the vicinity of haustoria cells were either devoid of fluorecence (**a, b**, star) or showed a destructured ER (**c, d**, asterisk), suggesting cell death. **e.** Using bright field microscopy, similar events were observed as destructuring of the vacuole (blue smear, asterisk) or light blue material filling the cell (star). These observations suggested rare occurrence of dying or dead cells in contact or in the vicinity of the haustorium.

Scale bar = 10 µm (**a, b)**; 50 µm (**c-e).**

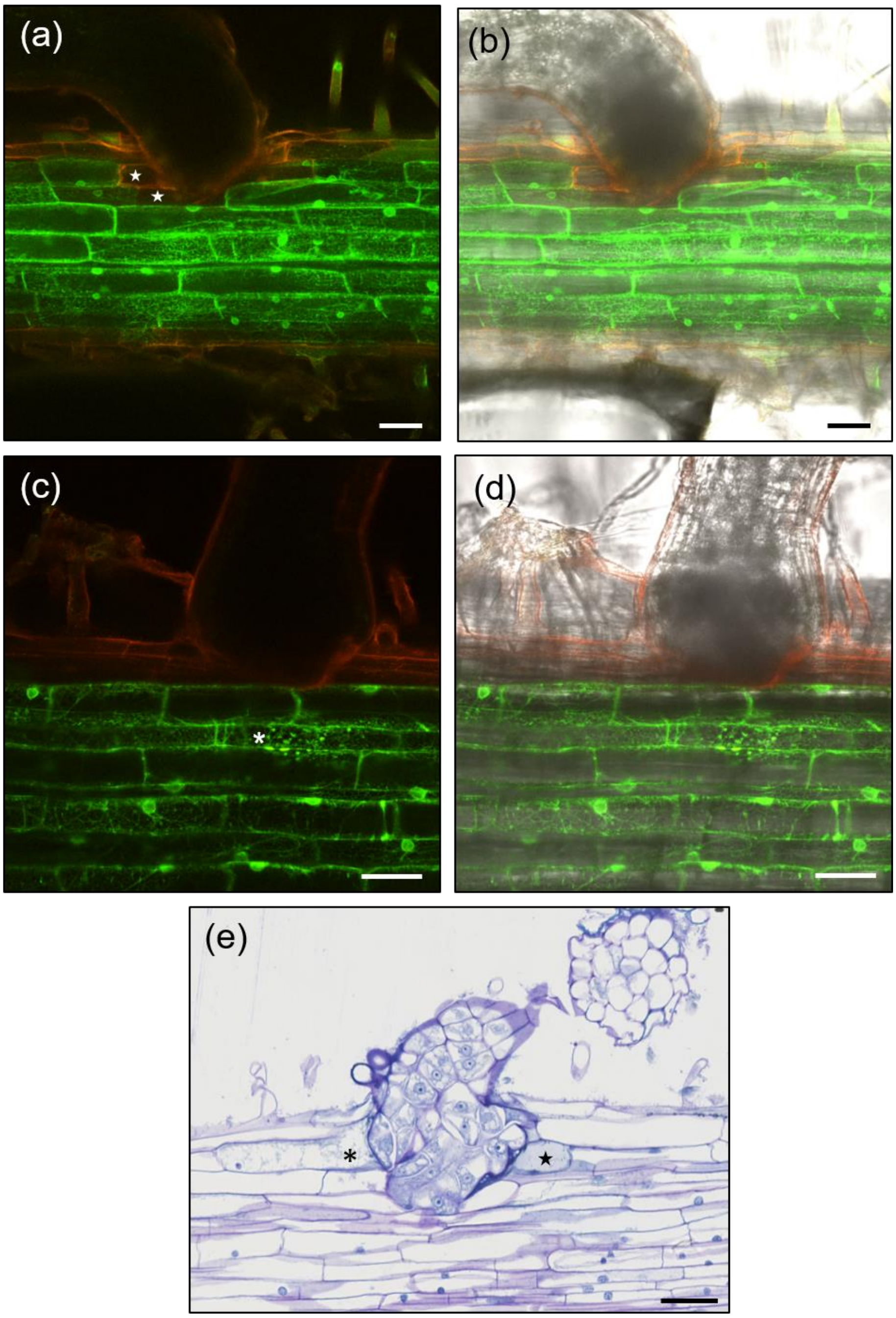

**Fig. S5. Multiple divisions in the host root at the site of broomrape intrusion.**

Lower magnification image of the section in **Fig. 1j,** showing a wider zone of the root. Cell division was observed around the intrusive cells (arrowheads). Anticlinal divisions affected the host cortex on both sides of the root (black square brackets). Periclinal divisions were visible in the pericycle (red square brackets).

Scale bar = 100 µm.

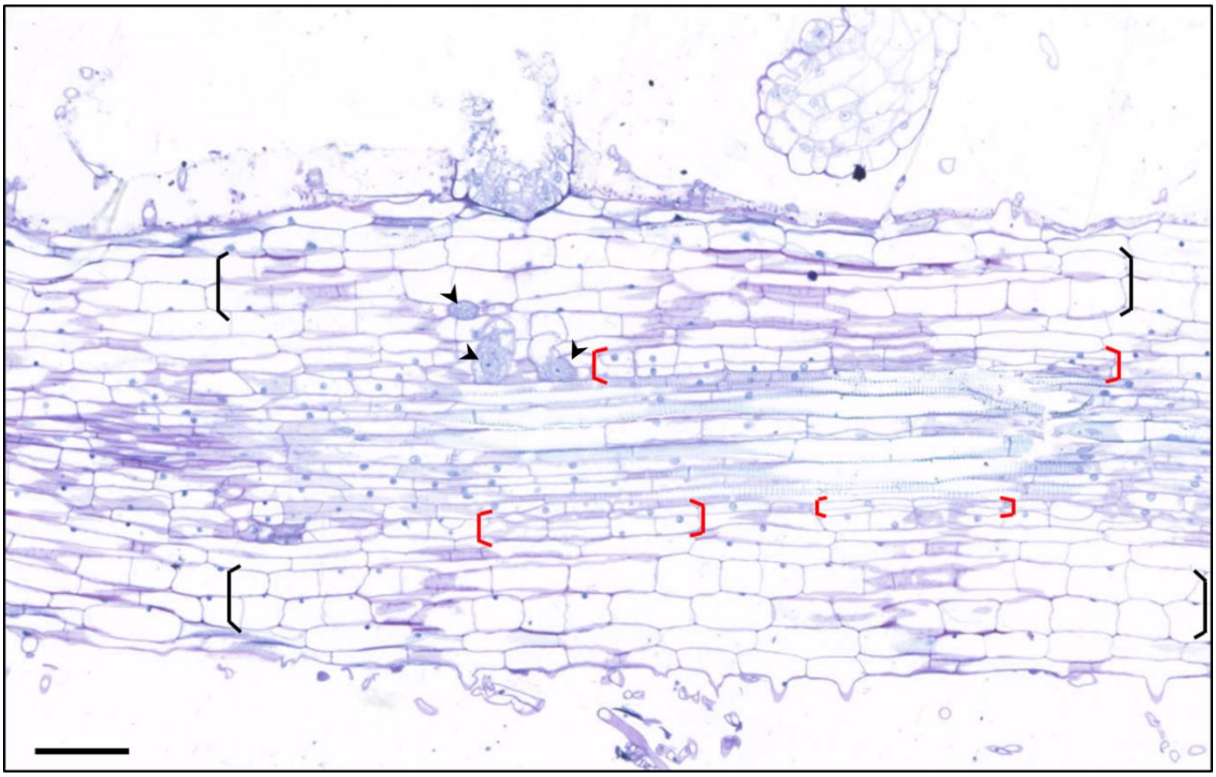

**Fig. S6. Transmission electron microscopy of a 7 dai attachment.**

The attachment shown in Fig. 2 was sectioned *c.* 9 µm further. **a.** General view of the attachment using large field microscopy. **b-d.** TEM.

**a.** Intrusive cells showed a dense cytoplasmic content and a big nucleus (arrow), containing a large nucleolus (dark blue). **b.** Assembly of 3 TEM pictures at the interface of the penetrating intrusive cells. The cell wall of the host cortical cell contacted by the tip of the haustorium was disaggregated (stars), potentially digested by parasitic pectolytic enzymes. By contrast the parasitic cell wall surrounding the intrusive cells (dotted line) was intact. The plasmalemma of the host cell showed convolution, suggesting active membrane synthesis, preparing the apoplastic compartment for subsequent accommodation of the haustorium. Electron dense granules were present along the host plasmalemma. **c, d**. One of the penetrated cortical cells, adjacent to the attachment, showed abnormal accumulation of granular material inside the vacuole. Electron microscopy of this cell indicated disruption of the tonoplast and plasmalemma as well as cytoplasm degradation, likely causes of cell death. Scale bar = 50 µm (**a**); 5 µm (**b**); 1 µm (**c-d**).

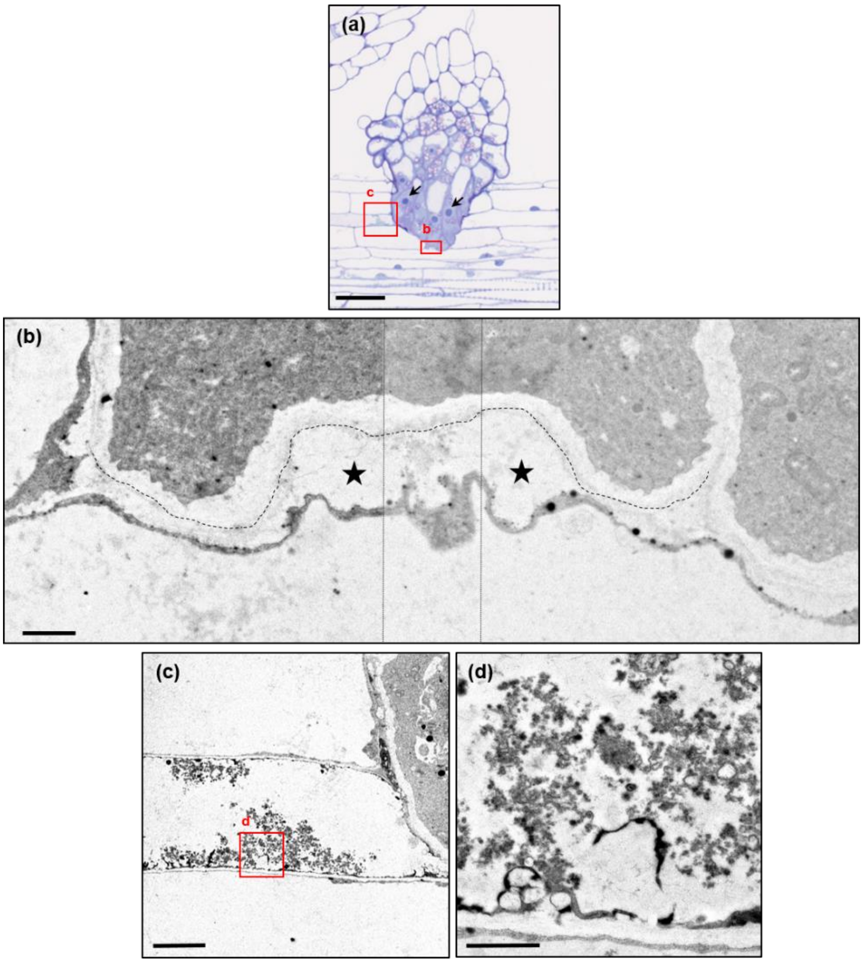

**Table S1. Efficiency of sunflower transformation *via A. rhizogenes.***

Four experiments of sunflower transformation were performed, using the p35S-GFP-ER construct (**Fig. S1**). Transgenic fluorescent roots were observed using a stereomicroscope equipped with a Long Pass GFP filter, which allowed to discriminate transgenic green roots from non-transformed orange auto-fluorescent roots. Plants were counted as transformed when at least one green fluorescent root had developed.

NA: data not available. In experiment No.1, composite plants were too developed and consequently transgenic roots were intermingled and difficult to distinguish from one another.

| experiment | number of plants | % of transformed plants | number of fluorescent root/ transformed plant | % of fluorescent root/ transformed plant |
|---|---|---|---|---|
| No.1 | 20 | 95 | NA | NA |
| No.2 | 7 | 86 | 4,6 | 63 |
| No.3 | 9 | 100 | 5,4 | 51 |
| No.4 | 10 | 100 | 4,9 | 49 |

**Table S2. Overview of observed sites.**

Twenty five attachments (4 to 8 dai) from inoculated wild-type (non transformed) sunflower plants were scored using large field microscopy: 21 sites of penetration and 4 sites without penetration yet. Thirty eight attachments (5 to 7 dai) were scored under confocal microscopy on inoculated transgenic composite plants: 21 sites of penetration and 17 sites without penetration yet. Sites are organized according to time of observation (dai) and stage of colonization (*i.e.* deepest sunflower root cell layer reached by the intrusive cells).

| | | microscopy | |
|---|---|---|---|
| Time | Stage of colonization | large field | confocal |
| 4 dai | no contact | 1 | |
| | epidermis | 1 | |
| 5 dai | contact but no penetration | | 3 |
| 6 dai | contact but no penetration | 3 | 10 |
| | epidermis | 1 | 1 |
| | 1st outer cortex | 1 | 3 |
| | 2nd or 3rd outer cortex | 1 | 8 |
| | inner cortex | 1 | 2 |
| | vessels | 1 | |
| 7 dai | contact but no penetration | | 4 |
| | epidermis | | 1 |
| | 1st outer cortex | 2 | 2 |
| | 2nd/ 3rd outer cortex | 4 | 3 |
| | inner cortex | 3 | 1* |
| | vessels | 1 | |
| 8 dai | 1st layer outer cortex | 2 | |
| | 2nd/ 3rd outer cortex | | |
| | inner cortex | 1 | |
| | endodermis | 1 | |
| | vessels | 1 | |
| | total number of sites | 25 | 38 |

*: inner cortex or deeper
